# Replicative fitness SARS-CoV-2 20I/501Y.V1 variant in a human reconstituted bronchial epithelium

**DOI:** 10.1101/2021.03.22.436427

**Authors:** Franck Touret, Léa Luciani, Cécile Baronti, Maxime Cochin, Jean-Sélim Driouich, Magali Gilles, Laurence Thirion, Antoine Nougairède, Xavier de Lamballerie

## Abstract

Since its emergence in 2019, circulating populations of the new coronavirus continuously acquired genetic diversity. At the end of 2020, a variant named 20I/501Y.V1 (lineage B.1.1.7) emerged and replaced other circulating strains in several regions. This phenomenon has been poorly associated to biological evidence that this variant and original strain exhibit different phenotypic characteristics. Here, we analyse the replication ability of this new variant in different cellular models using for comparison an ancestral D614G European strain (lineage B1). Results from comparative replication kinetics experiments *in vitro* and in a human reconstituted bronchial epithelium showed no difference. However, when both viruses were put in competition in a human reconstituted bronchial epithelium, the 20I/501Y.V1 variant outcompeted the ancestral strain. Altogether, these findings demonstrate that this new variant replicates more efficiently and could contribute to better understand the progressive replacement of circulating strains by the SARS-CoV-2 20I/501Y.V1 variant.

**Importance:** The emergence of several SARS-CoV-2 variants raised numerous questions concerning the future course of the pandemic. We are currently observing a replacement of the circulating viruses by the variant from the United Kingdom known as 20I/501Y.V1 from B.1.1.7 lineage but there is little biological evidence that this new variant exhibit a different phenotype. In the present study, we used different cellular models to assess the replication ability of the 20I/501Y.V1 variant. Our results showed that this variant replicate more efficiently in a human reconstituted bronchial epithelium, which may explain why it spreads so rapidly in human populations.

## Observation

The new coronavirus SARS-Cov-2 emerged in China by the end of 2019 and rapidly spread worldwide. In a few months, D614G spike mutation was rapidly fixed in almost all SARS-CoV-2 circulating populations without evidence for higher COVID-19 mortality or clinical severity (1).It is still being debated whether it is due to a random founder effect (1) or, more probably, if the mutation enhances viral loads in the upper respiratory tract increasing the infectivity and stability of virions (2, 3).

In September 2020, emerged in the United Kingdom a variant named 20I/501Y.V1 from lineage B.1.1.7 (initially named VOC 2 2020212/01). It spread rapidly and is becoming dominant in Western Europe (4) and the USA (5). There is consistent epidemiological evidence that this so-called ‘UK variant’ is more efficiently transmitted than the pre-existing European strains, in particular in young patients. However, this has been poorly associated to biological evidence that the UK and original European variants exhibit different phenotypic characteristics.

Here, we present a comprehensive analysis of the replication ability *in vitro* and *ex vivo* of the 20I/501Y.V1 variant (strain UVE/SARS-CoV-2/2021/FR/7b isolated in February 2021 in Marseille, France; GISAID accession nb. EPI_ISL_918165), using for comparison the lineage B.1 BavPat D614G strain that circulated in Europe in February/March 2020.

The first experiments, performed in two cell lines commonly used for SARS Cov-2 culture, VeroE6/TMPRSS2 and Caco-2, revealed highly similar replication kinetics (Fig 1A-B). We then assessed the replicative fitness of both strains using a previously described model of reconstituted human airway epithelium (HAE) of bronchial origin (6). Following the inoculation of the epithelia through their apical side at a multiplicity of infection (MOI) of 0.1 in order to mimic the natural route of infection, we monitored the excretion of new virions at the apical side between 2 and 4 days post-infection (dpi) and measured the intracellular viral RNA yields at 4 dpi. Infectious titres (Fig.1D) and viral RNA yields (Fig.1E) at the apical side at 3 and 4 dpi, as well as intracellular viral RNA yields at 4 dpi (Fig. 1G), were slightly higher for the 20I/501Y.V1 variant. However, differences were not significant and estimated relative virion infectivity (*i*.*e*., the ratio of the number of infectious particles over the number of viral RNA) were similar for both viruses at all sampling times (Fig1. F). Altogether, these results are in line with our findings in common cell lines and with a recent report (7).

**Figure 1.**
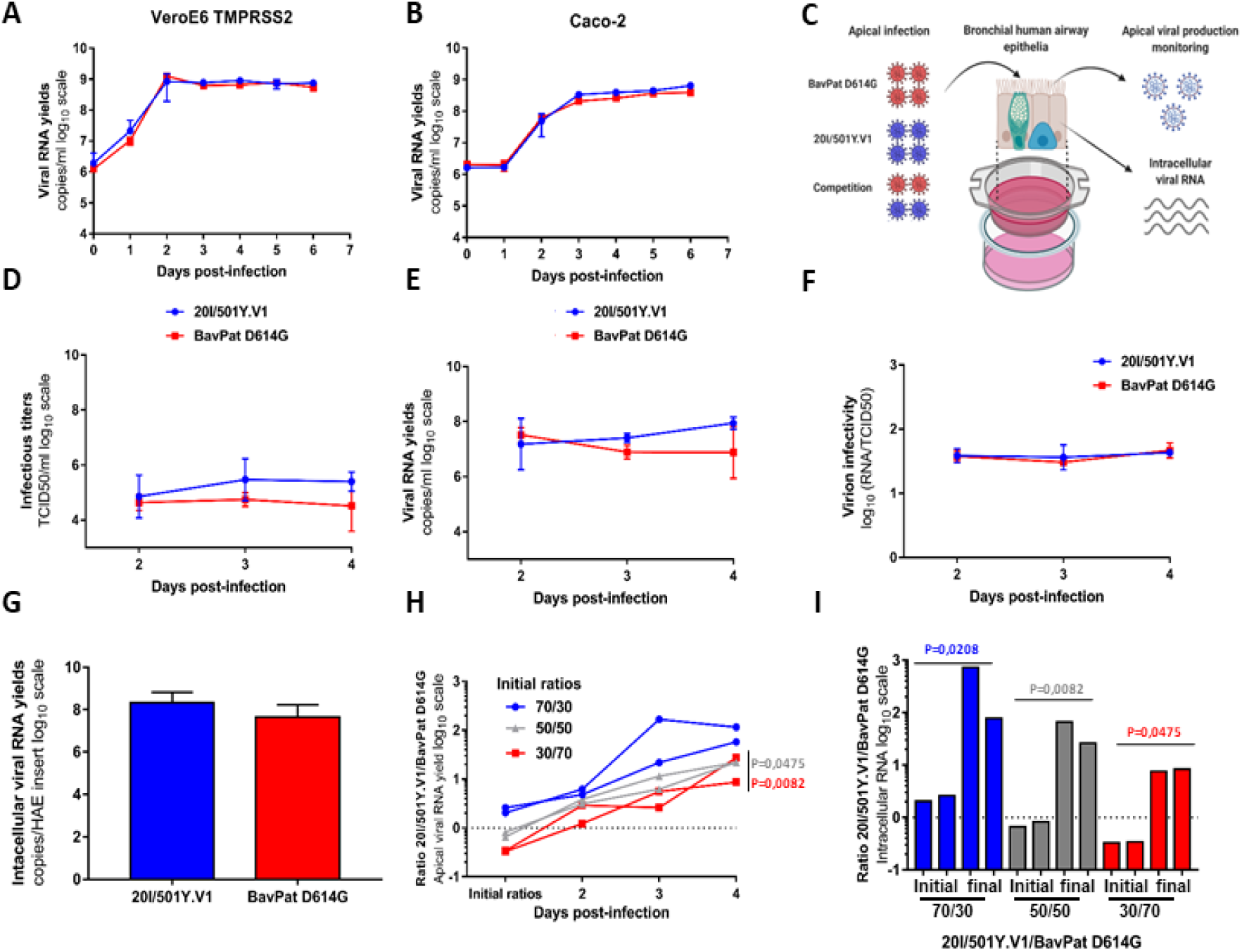
*In vitro* and *ex vivo* replication ability of a 20I/501Y.V1 variant in comparison with a lineage B.1 D614G strain. (A-B) Replication kinetics in VeroE6 TMPRSS2 (A) and Caco-2 (B) cells. Viral replication was assessed using a RT-qPCR assay. (C) Graphical representation of experiments with reconstituted human airway epithelium (HAE) of bronchial origin. (D-E) Kinetics of virus excretion at the apical side of the epithelium measured using a TCID_50_ assay (D) and a RT-qPCR assay (E). (F) Estimation of virion infectivities (*i*.*e*., the ratio of the number of infectious particles over the number of viral RNA). (G) Intracellular viral RNA yields measured at 4 dpi using a RT-qPCR assay. (A-G) Data represent mean±SD of a triplicate. No statistical difference was observed between both viral strains (p>0.05; unpaired Mann-Whitney test). (H) Follow-up of the 20I/501Y.V1/BavPat D614G ratios at the apical side. Each line represents results from an HAE insert. (I) Individual 20I/501Y.V1/BavPat D614G ratios estimated from intracellular viral RNAs at 4dpi (I). (H-I) p-values were determined against the initial ratios using Kruskal-Wallis test followed by uncorrected Dunn’s post-hoc analysis. The graphical representation was created with BioRENDER.

Based on these results, we performed competition experiments, which have been previously demonstrated to be effective to detect moderate replicative fitness differences(8). Accordingly, we inoculated epithelia with both viruses simultaneously as described above, sampled the apical side between 2 and 4 dpi and extracted intracellular viral RNA yields at 4 dpi. Three infection inoculum ratios (20I/501Y.V1/BavPat D614G: 70/30, 50/50 and 30/70) were used. Using two specific RT-qPCR assays, we estimated the proportion of each viral genome in the viral population (expressed as 20I/501Y.V1/BavPat D614G ratio in Figure 1H-I). Results revealed that regardless the initial ratio, a similar pattern was observed in which Bavpat D614G was outcompeted by the 20I/501Y.V1 variant: all 20I/501Y.V1/BavPat D614G ratio values estimated from apical side washes were above 1 and over 57, 22 and 8 at 4 dpi for epithelia inoculated for initial ratios 70/30, 50/50 and 30/70 respectively (Fig1. H). Notaby, 20I/501Y.V1/BavPat D614G ratios measured at 4 dpi were significantly higher than initial 50/50 and 30/70 inoculum ratios (p= 0.0475 and p=0.0082 respectively; Kruskal-Wallis test with uncorrected Dunn’s post-hoc analysis). Similar result were observed when estimating the 20I/501Y.V1/BavPat D614G ratios from intracellular viral RNAs (Fig1. I): 20I/501Y.V1/BavPat D614G ratios measured at 4 dpi were significantly higher than initial 50/50 and 30/70 inoculum ratios (p= 0.0208, p=0.0082 and p=0.0475 with 70/30, 50/50 and 30/70 inoculum ratios respectively; Kruskal-Wallis test with uncorrected Dunn’s post-hoc analysis).

Our results demonstrated that the 20I/501Y.V1 is more fit than the BavPat D614G in a reconstituted bronchial human epithelium. This may be explained by the presence of the N501Y mutation in the receptor binding domain (RBD) of the spike protein which enhances viral particle binding to the ACE2 receptor (9). This may translate into a fitness advantage as demonstrated in a recent study with engineered viral strains (10). Similar observations have been made with the D614G mutation, where the new G614 strains overcame the D614 original strains when put in competition (2). Altogether, these findings could contribute to better understand the progressive replacement of circulating strains by the SARS-CoV-2 20I/501Y.V1 variant (11).

## Acknowledgments

We thank Pr C Drosten for providing the SARS-CoV-2 BavPat strain through EVA GLOBAL. This work was supported by Inserm through the REACTing (REsearch and ACTion targeting emerging infectious diseases) initiative. This work was supported by the European Virus Archive Global (EVA GLOBAL) funded by the European Union’s Horizon 2020 research and innovation programme under grant agreement No 871029. This work was supported by the Fondation de France “call FLASH COVID-19”, project TAMAC.

