## Supplemental for "Replicative fitness SARS-CoV-2 20I/501Y.V1 variant in a human reconstituted bronchial epithelium"

\# Co-last authors

Keywords: SARS-CoV-2; variant; replicative fitness; *ex vivo*

Supplemental materials and methods

Cell lines

VeroE6 (ATCC CRL-1586) ), VeroE6/TMPRSS2 (NIBSC 100978) and Caco-2 (ATCC HTB-37) cells were grown in minimal essential medium (MEM, Life Technologies), supplemented with 5% heat-inactivated fetal calf serum (FCS; Life Technologies), 1% penicillin/streptomycin (PS, 5000U.mL^−1^ and 5000µg.mL^−1^ respectively; Life Technologies), 1 % non-essential amino acids (Life Technologies) and L-Glutamine (Life Technologies), at 37°C with 5% CO_2_.

Human airway epithelia (HAE)

Mucilair™ HAE reconstituted from human primary cells of bronchial biopsies were purchased from Epithelix SARL (Geneva, Switzerland). The bronchial epithelium was maintained in air-liquid interface with specific media purchased from Epithelix. Epithelium derived from a 17-year-old donor Hispanic male with no existing pathologies reported or detected. All samples purchased from Epithelix SARL have been obtained with informed consent. These studies were conducted according to the declaration of Helsinki on biomedical research (Hong Kong amendment, 1989), and received approval from local ethics committee.

Virus strain

SARS-CoV-2 strain BavPat1 D614G was obtained from Pr. C. Drosten through EVA GLOBAL (https://www.european-virus-archive.com/). SARS-CoV-2 201/501YV.1 was isolated from a 18 years-old patient (full-length genome sequence deposited on GISAID : EPI_ISL_918165) and is available through EVA GLOBAL (UVE/SARS-CoV-2/2021/FR/7b; lineage B 1. 1 .7, ex UK; <https://www.european-virus-archive.com/virus/sars-cov-2-uvesars-cov-22021fr7b-lineage-b-1-1-7-ex-uk>.

To prepare the virus working stock, a 25cm^2^ culture flask of confluent VeroE6 cells growing with MEM medium supplemented with 2.5% FCS was inoculated at multiplicity of infection (MOI) of 0.001. Cell supernatant medium was harvested at the peak of replication and supplemented with 25mM HEPES (Sigma-Aldrich) before being stored frozen in aliquots at -80°C. All experiments with infectious virus were conducted in a biosafety level 3 laboratory.

Replication kinetics in Caco-2 and VeroE6/TMPRSS2

One day prior infection 5×10^4^ VeroE6 TMPRSS2 or 7×10^4^ Caco-2 cells were seeded in 96 well plates with 100µL assay medium (containing 2.5% FCS) per well. The next day, cells were infected at a MOI of 0.01. Every day, 100µl of cell supernatant, in triplicate, was collected in a S-Block (Qiagen) previously loaded with VXL lysis buffer containing proteinase K and RNA for further analysis.

HAE experiments

After a gentle wash with pre-warmed OPTI-MEM medium ( Life technologies), epithelia were infected with SARS-COV-2 or a mix of both viruses (competition experiments) on the apical side using a MOI of 0.1 as previously described (Pizzorno et al., 2020). For competition assay, ratios were calculated based on TCID_50_ measurement. Even in the competition experiment the final MOI was 0.1. At day 1, the apical side of the epithelium was washed with pre-warmed OPTI-MEM in order to eliminate the viral inoculum. Samples were collected at the apical side by washing with 200µL of pre-warmed OptiMEM medium: 100µL was collected in S-Block (Qiagen) previously loaded with VXL lysis buffer containing proteinase K and RNA carrier for RNA extraction; the remaining quantity was used to perform a TCID_50_ assay_._

At day 4 post-infection, each epithelium were lysed with ATL (Qiagen) and load into a S-Block (Qiagen) for extraction of intracellular RNAs.

Quantification of the viral genome by real-time RT-qPCR

To avoid contamination, all experiments were conducted in a molecular biology laboratory that was specifically designed for clinical diagnosis, and which includes separate laboratories for each step of the procedure. RNA extraction was performed using the Qiacube HT automat and the QIAamp 96 DNA kit HT following manufacturer instructions. Total intracellular RNA of each well was extracted using RNeasy 96 HT kit (Qiagen) following manufacturer’s instructions. Viral RNA was quantified by real-time RT-qPCR (GoTaq 1-step qRt-PCR, Promega) using 3.8µL of extracted RNA and 6.2µL of RT-qPCR mix and standard fast cycling parameters, *i.e.*, 10min at 50°C, 2 min at 95°C, and 40 amplification cycles (95°C for 3 sec followed by 30sec at 60°C). Quantification was provided by four 2 log serial dilutions of an appropriate T7-generated synthetic RNA standard of known quantities (10^2^ to 10^8^ copies/reaction). RT-qPCR reactions were performed on QuantStudio 12K Flex Real-Time PCR System (Applied Biosystems) and analyzed using QuantStudio 12K Flex Applied Biosystems software v1.2.3. Primers and probe sequences, which target SARS-CoV-2 N gene, were: Fw: GGCCGCAAATTGCACAAT ; Rev : CCAATGCGCGACATTCC; Probe: FAM-CCCCCAGCGCTTCAGCGTTCT-BHQ1.

Specific quantification of 20I/501Y.V1 and BavPat D614G strains by real-time RT-qPCR

Two specific rea-time RT-qPCR assays (each specifically detecting one of the competing viruses) were used to determine the proportion of each viral genome in all samples. Prior to PCR amplification, RNA extraction was performed as described above (Qiacube HT automat and the QIAamp 96 DNA kit HT).

RT-qPCR (SARS-CoV- 2) were performed with the Kit SuperScript III Platinum One-Step qRT-PCR (Kits SuperScript™ III Platinum™ One-Step qRT-PCR  Kit, universal Invitrogen) using 2.5µL of RNA and 7.5µL of RT qPCR mix and standard fast cycling parameters, i.e., 15min at 50°C, 2 min at 95°C, and 40 amplification cycles (95°C for 15 sec followed by 45 sec at 55°C). Quantification was provided by four 2 log serial dilutions of an appropriate T7-generated synthetic RNA standard of known quantities (107 to 10^3^ copies/reaction). RT-qPCR reactions were performed on QuantStudio 12K Flex Real-Time PCR System (Applied Biosystems) and analyzed using QuantStudio 12K Flex Applied Biosystems software v1.2.3. Primers and probes sequences used to detect SARS-CoV-2 : NSP6 Fwd: 5'- CAT GGT TGG ATA TGG TTG-3' Rev: 5'- GAT GCA TAC ATA ACA CAG-3' Probe that specifically detect the BavPat D614G : 5'- : FAM-GTC TGG TTT TAA BHQ1-3'; Probe that specifically detect the 201/501YV.1  VIC-TAG TTT GAA GCT-BHQ1-3'.

Tissue-culture infectious dose 50 (TCID50) assay

To determine infectious titers, 96-well culture plates containing confluent VeroE6 cells were inoculated with 100μL per well of ten-fold dilutions serial dilutions of each sample in sextuplicate. Plates were incubated for 5 days and then read for the absence or presence of cytopathic effect in each well. Infectious titers were estimated using the method described by Reed & Muench (Reed and Muench, 1938) and expressed as TCID_50_/ml.

Statistical tests are described in the figure legend. All data obtained were analyzed using GraphPad Prism 7 software (Graphpad software).
